# OligoMiner: A rapid, flexible environment for the design of genome-scale oligonucleotide *in situ* hybridization probes

**DOI:** 10.1101/171504

**Authors:** Brian J. Beliveau, Jocelyn Y. Kishi, Guy Nir, Hiroshi M. Sasaki, Sinem K. Saka, Son C. Nguyen, Chao-ting Wu, Peng Yin

**Author notes:** Present address: Department of Genetics, University of Pennsylvania, Philadelphia, PA 19104. To whom correspondence may be addressed. Mail: 3 Blackfan Circle CLS Room 543, Boston, MA, 02115.

## Abstract

Oligonucleotide (oligo)-based fluorescence in situ hybridization (FISH) has emerged as an important tool for the study of chromosome organization and gene expression and has been empowered by the commercial availability of highly complex pools of oligos. However, a dedicated bioinformatic design utility has yet to be created specifically for the purpose of identifying optimal oligo FISH probe sequences on the genome-wide scale. Here, we introduce OligoMiner, a rapid and robust computational pipeline for the genome-scale design of oligo FISH probes that affords the scientist exact control over the parameters of each probe. Our streamlined method uses standard bioinformatic file formats, allowing users to seamlessly integrate existing and new utilities into the pipeline as desired, and introduces a novel method for evaluating the specificity of each probe molecule that connects simulated hybridization energetics to rapidly generated sequence alignments using supervised learning. We demonstrate the scalability of our approach by performing genome-scale probe discovery in numerous model organism genomes and showcase the performance of the resulting probes with both diffraction-limited and single-molecule super-resolution imaging of chromosomal and RNA targets. We anticipate this pipeline will make the FISH probe design process much more accessible and will more broadly facilitate the design of pools of hybridization probes for a variety of applications.

## Introduction

Fluorescence *in situ* hybridization (FISH) is a powerful single-cell technique that harnesses the specificity afforded by Watson-Crick base pairing to reveal the abundance and positioning of cellular RNA and DNA molecules in fixed samples. Originally introduced as a radioactive *in situ* hybridization method in the late 1960s (1–3), FISH has undergone a series of optimizations that have improved its detection efficiency and sensitivity (4–7). Many of these refinements have centered on the preparation and labeling of the probe material, which traditionally has been derived from cellular DNA or RNA, and include the introduction of the nick translation method that increases the specific activity of labeling (8, 9) and the development of suppressive hybridization techniques that limit background originating from repetitive sequences contained in many probes (10).

More recently, advances in DNA synthesis technology have afforded researchers the opportunity to construct FISH probes entirely from synthetic oligonucleotides (oligos). Oligo probes offer many potential advantages, as they can be selected to have specific thermodynamic properties, engineered to avoid repetitive sequences, designed against any sequenced genome, and endowed with many different types and densities of labels. While the use of oligo probes was initially restricted to the interrogation of multi-copy targets such as repetitive DNA (11–13) and messenger RNA (14–16) using one to a few dozen oligo probes, the recent development of oligo libraries produced by massively parallel array synthesis (17) has empowered a new generation of FISH technologies able to target single-copy chromosomal regions with highly complex libraries of hundreds to many thousands of oligo probes (18–20).

We have previously introduced Oligopaints, a method for the generation of highly efficient probes for both RNA FISH and DNA FISH from libraries composed of dozens to many thousands of unique oligo species (20). A key feature of Oligopaints is their programmability, wherein the genomic and non-genomic sequences that compose each probe oligo can be specified precisely. This fine level of control has enabled several important technical advances in FISH imaging, including the single-molecule super-resolution imaging of chromosome structure at non-repetitive targets (21, 22), the development of probes that can distinguish genomically unique regions of homologous chromosomes (21), and the introduction of a method able to label dozens of chromosomal loci (23). The general programmability of oligo FISH probes has also enabled the creation of related methods that utilize aspects of the Oligopaints approach to enable the highly multiplexed visualization of dozens to >1000 distinct mRNA species in the same sample (24, 25).

Despite the rapid maturation of new FISH technologies reliant upon oligo probes, comparatively little progress has been made in the development of computational tools to facilitate the design of these oligos. While computational utilities exist to create small numbers of oligo probes against targets such as bacterial ribosomal RNA (26, 27) and to design large pools of oligo pairs such as PCR primers (28–31) or padlock probes (32, 33), to our knowledge no bioinformatic utility has been created for the explicit purpose of designing oligo hybridization probes at the genome-wide scale. Consequently, older utilities such as the microarray design program OligoArray (34) have been repurposed to facilitate probe design. Although OligoArray has produced effective oligo FISH probes (20–24), it can only provide limited throughput, with large genomes such as those of human and mouse taking 1–2 months of continuous cluster computing to mine with a single set of parameters (20) and smaller regions still requiring hours of cluster computing to complete. Additionally, OligoArray offers the user a limited amount of control over the probe discovery process, as users interact only with a compiled executable Java Archive file and cannot change the nature or order of steps taken or the values of many of parameters used for thermodynamic calculations and specificity checking.

Here, we introduce OligoMiner, a rapid and flexible genome-scale design environment for oligo hybridization probes. The modular, open source OligoMiner pipeline is written in Python and Biopython (35) and uses standard bioinformatic file formats at each step in the probe mining process, greatly simplifying probe discovery. Additionally, OligoMiner introduces a novel method of assessing probe specificity that employs supervised machine learning to predict thermodynamic behavior from genome-scale sequence alignment information. The OligoMiner pipeline can readily be deployed on any sequenced genome and can mine the entirety of the human genome in minutes to hours and smaller <10 Mb regions in mere minutes on a standard desktop or laptop computer, greatly reducing the time and computational resource cost of probe discovery. We also demonstrate the effectiveness of probes produced by our approach with both conventional and single-molecule super-resolution microscopy.

## Results

### Identification of Candidate Probes

The OligoMiner workflow begins with a FASTA-formatted input file (36) containing the genomic sequence to be searched for probes, which can be masked by a program such as RepeatMasker (37) to exclude regions containing repetitive elements. This input file is first passed to the blockParse script (Fig. 1*A* and Fig. S1), which screens for prohibited sequences such as homopolymeric runs and ‘N’ bases and allows users to specify allowable ranges of probe length, percent G+C content (GC%), and adjusted melting temperature (*T*_*m*_) calculated using nearest neighbor thermodynamics (38). Candidate probe sequences passing all checks are outputted in FASTQ format (39) to facilitate input into next generation sequencing (NGS) alignment programs such as Bowtie/Bowtie2 (40, 41) and BWA (42), which can be used to assess off-target potential. Importantly, these NGS alignment programs are optimized for the extremely rapid alignment of millions of short sequences to a reference genome in parallel, thus allowing the specificity check step of the pipeline to proceed much more quickly than approaches like OligoArray that use BLAST (43) in serial.

**Fig. 1.**
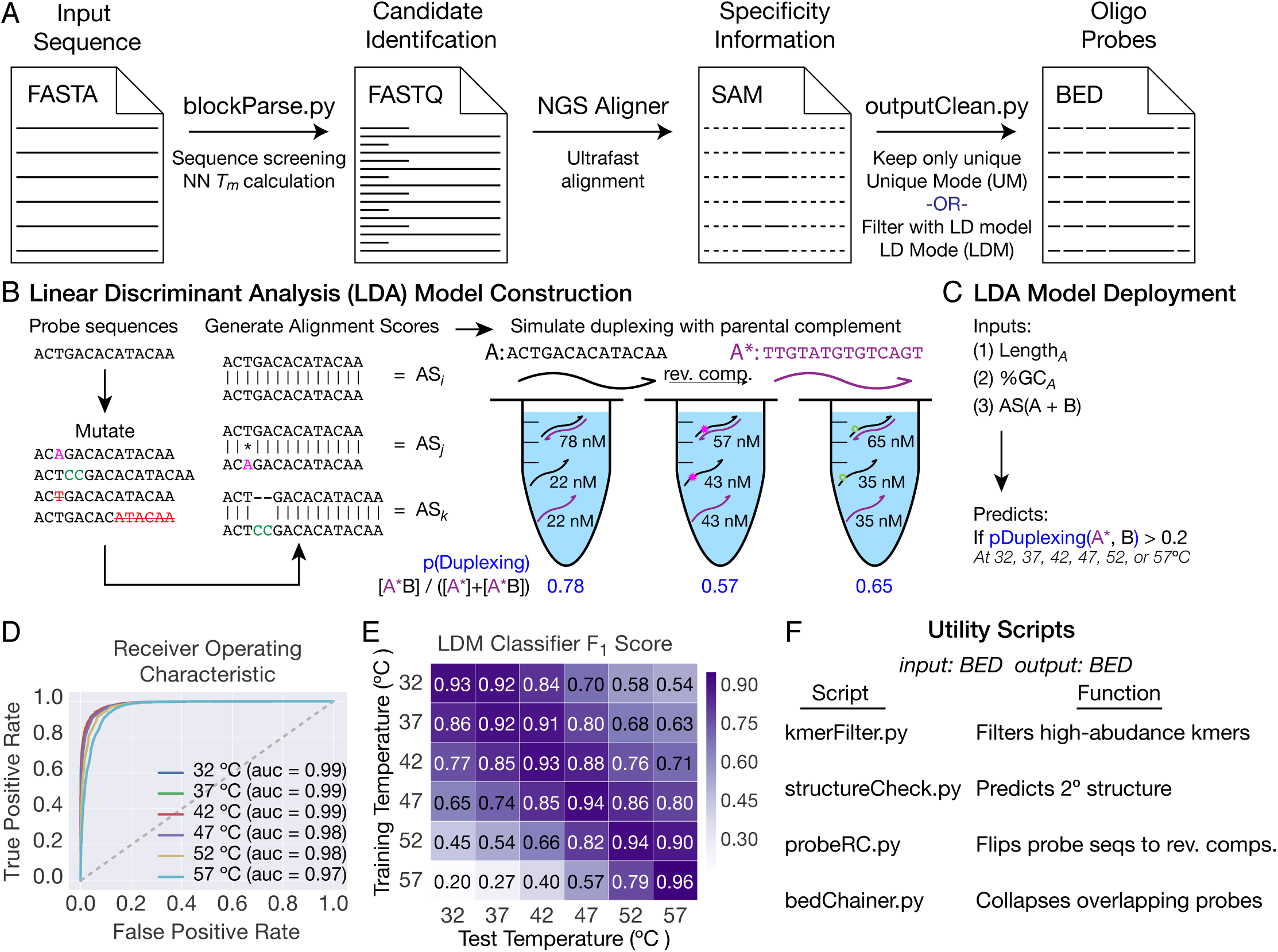
Implementing OligoMiner. (*A*) Schematic overview of the OligoMiner pipeline. (*B* and *C*) Schematic overviews of Linear Discriminant Analysis (LDA) model (*B*) creation and (*C*) implementation. (*D*) Receiver Operating Characteristic curves for each temperature-specific LDA model. ‘auc’ = area under the curve. (*E*) Heat map showing the support-weighted F_1_ score for each temperature-specific LDA model when tested against validation data simulated at each of the six indicated temperatures. (*F*) Description of utility scripts also developed as part of OligoMiner.

### Predicting Probe Specificity

Ultrafast alignment programs can provide a wealth of information about the relatedness of a given input sequence to other sequences present in a genome assembly. OligoMiner allows users to evaluate probe specificity using two distinct approaches, in either case using the script outputClean to process the Sequence Alignment/Map (SAM) file (44) produced by the alignment program and outputting Browser Extendable Data (BED) format (45) files; BED files are designed for visualizing sequence features in genome browsers and are fully compatible with our previously described tools that facilitate the design and ordering of Oligopaint probe libraries (20) (http://genetics.med.harvard.edu/oligopaints) and utilities such as BEDTools (46) (Fig. 1*A* and Fig. S2). The first approach, termed ‘Unique Mode’ (UM), uses the number of reported alignments to differentiate between candidate probes predicted to only have one genomic target from those with multiple potential binding sites; candidates with more than one reported alignment or that fail to align are filtered, while candidate probes that align uniquely are passed to the output. ‘Unique Mode’ thus enables users to experiment with different groups of alignment parameters in order to find an optimal combination for a given application.

Ideally, the thermodynamics of hybridization between a candidate probe and potential off-target sites would be modeled *in silico* and employed as a means of identifying probe oligos likely to only bind their intended targets in a given set of reaction conditions. While powerful utilities such as NUPACK (47–49) are capable of performing such simulations, the limited throughput of these programs renders a direct thermodynamic approach impractical for genome-scale probe design. Yet, we hypothesized that features in rapidly calculated data such as alignment scores may be predictive of thermodynamic behavior and could thus serve as a proxy for the information that would be produced by thermodynamic simulations. Inspired by this idea, we first selected 800 ‘probe’ sequences identified by blockParse in the human hg38 assembly that represented three commonly used probe length ranges (26–32, 35–41, 40–46 nt) (*Methods*). In order to simulate the types of binding sites that these ‘probes’ might encounter *in situ* during a FISH experiment in a complex genome, we next generated 406,014 variant versions of the ‘probe’ sequences *in silico* that each contained one or more point mutation, insertion, deletion, or large truncation, creating in combination with the 800 ‘probe’ sequences a pool of 406,814 ‘target sites’ (*Methods*) (Fig. 1*B*). We then aligned each ‘probe’ to its corresponding ‘target sites’ in pairwise alignments using Bowtie2 with ultrasensitive settings (*Methods*), generating a set of 406,814 alignment scores (Fig. 1*B*). In parallel, we also computed the probability of a duplex forming between each ‘probe’ and each of its corresponding ‘target sites’ in FISH conditions (2X SSC, 50% formamide at 32, 37, 42, 47, 52, or 57 °C) in pairwise test tube simulations using NUPACK (*Methods*) (Fig. 1*B*).

In order to connect our alignment scores and duplexing probabilities, we next performed supervised machine learning using linear discriminant analysis (LDA) on 60% of the combined datasets with scikit-learn (50). Specifically, we built six temperature-specific LDA models that predict whether the duplexing probability of a ‘probe’ – ‘target site’ pair will be above a threshold level of 0.2 (i.e. <5-fold weaker than a fully paired duplex) given the length and GC% of the ‘probe’ sequence and the score of the alignment of the two sequences (*Methods*) (Fig. 1*C*). We tested these LDA models on the remaining 40% of the data and found that all six performed exceptionally well, with each producing areas under receiver operating characteristic curves of ≥0.97 (Fig. 1*D*) and support-weighted F_1_ scores ≥0.92 (Fig. 1*D* and Fig. S3). Notably, all six models also performed strongly when tested against data simulated at hybridization temperatures 5°C higher or lower than the training temperature (support-weighted F_1_ score range 0.79–0.92, mean 0.86; Fig. 1*E*), indicating that the models are all capable of predicting duplexing behavior over a relatively broad range of reaction conditions. Collectively, our data argue that the LDA model identifies potentially problematic ‘probe’ – ‘target site’ interactions (i.e. those with a probability of duplexing >0.2) effectively as well as the much slower thermodynamic simulations. We have integrated the six LDA models into outputClean to create the second specificity evaluating approach, ‘LDA Mode’ (LDM): candidate probes are first aligned to the reference genome of interest using the same Bowtie2 scoring settings used to construct the LDA models (*Methods*), and the resulting SAM file is processed by a selected temperature-specific LDA model such that candidate probes predicted to have more than one thermodynamically relevant target site (probability of duplexing >0.2) are filtered (Fig. 1*A* and Fig. S2).

### Post-Processing Functionalities

We have written a series of utility scripts to augment the core OligoMiner pipeline (Fig. 1*F*). These utility scripts accept and return BED files, making them compatible with both output files created by outputClean (Fig. 1*A*) and files created by the previous Oligopaint probe discovery method (20) and adding additional functionalities. For instance, kmerFilter enables the user to perform another layer of specificity checking by calling Jellyfish (51) to screen probe sequences for the presence of high abundance k-mers (e.g. 16mers or 18mers) that may be missed by alignment programs due to their short lengths and could lead to off-target binding (52, 53).

Users can also identify and filter probe sequences predicted to adopt secondary structures in a given set of experimental conditions using structureCheck, which depends on NUPACK. Several additional tools facilitate the processing of probe files for specific applications, including the conversion of probe sequences to their reverse complements by probeRC for strand-specific DNA or RNA FISH and the collapsing of overlapping probes by bedChainer for the design of high-density probe sets. Finally, we have created additional modularity with pair of scripts ‘fastqToBed’ and ‘bedToFastq’ that allow users to convert between the BED and FASTQ format files.

### Rapid Genome-Scale Probe Discovery

In order to assess the scalability of OligoMiner, we performed genome-wide probe discovery in the human hg38 genome assembly. We first developed three sets of input parameters spanning a range of commonly used probe lengths and experimental conditions: a ‘coverage’ set designed to maximize the number of probes discovered (26–32 nt length, 37°C hybridization), a ‘stringent’ set designed to maximize probe binding affinity and thus permit stringent hybridization and washing conditions (40–46 nt, 47°C hybridization), and a ‘balance’ set that seeks to compromise between coverage and binding affinity (35–41 nt, 42°C hybridization) (Fig. *2A*). We next deployed OligoMiner using these parameter settings in both UM and LDM, in both cases using Bowtie2 for the alignment step and also including the optional kmerFilter specificity check (*Methods*). Excitingly, both approaches were able to mine the entire hg38 assembly very rapidly using all three parameter sets, with UM averaging a rate of 1.70 Mb/minute and a total time of 97 minutes per chromosome across all three parameter settings (Fig. 2 *B* and *C*) and LDM averaging a similar rate of 1.48 Mb/minute and a total time of 104 minutes per chromosome (Fig. 2 *C* and *D*). These rates support mining the entire human genome in as little as 24–48 hours if each chromosome was run in serial on a laptop or desktop computer and tens of minutes if parallel computing (e.g. ∼100–400 simultaneous jobs) was instead employed, in either case achieving a dramatic increase in speed from the 1–2 months of parallel computing needed in our previous approach (20).

**Fig. 2.**
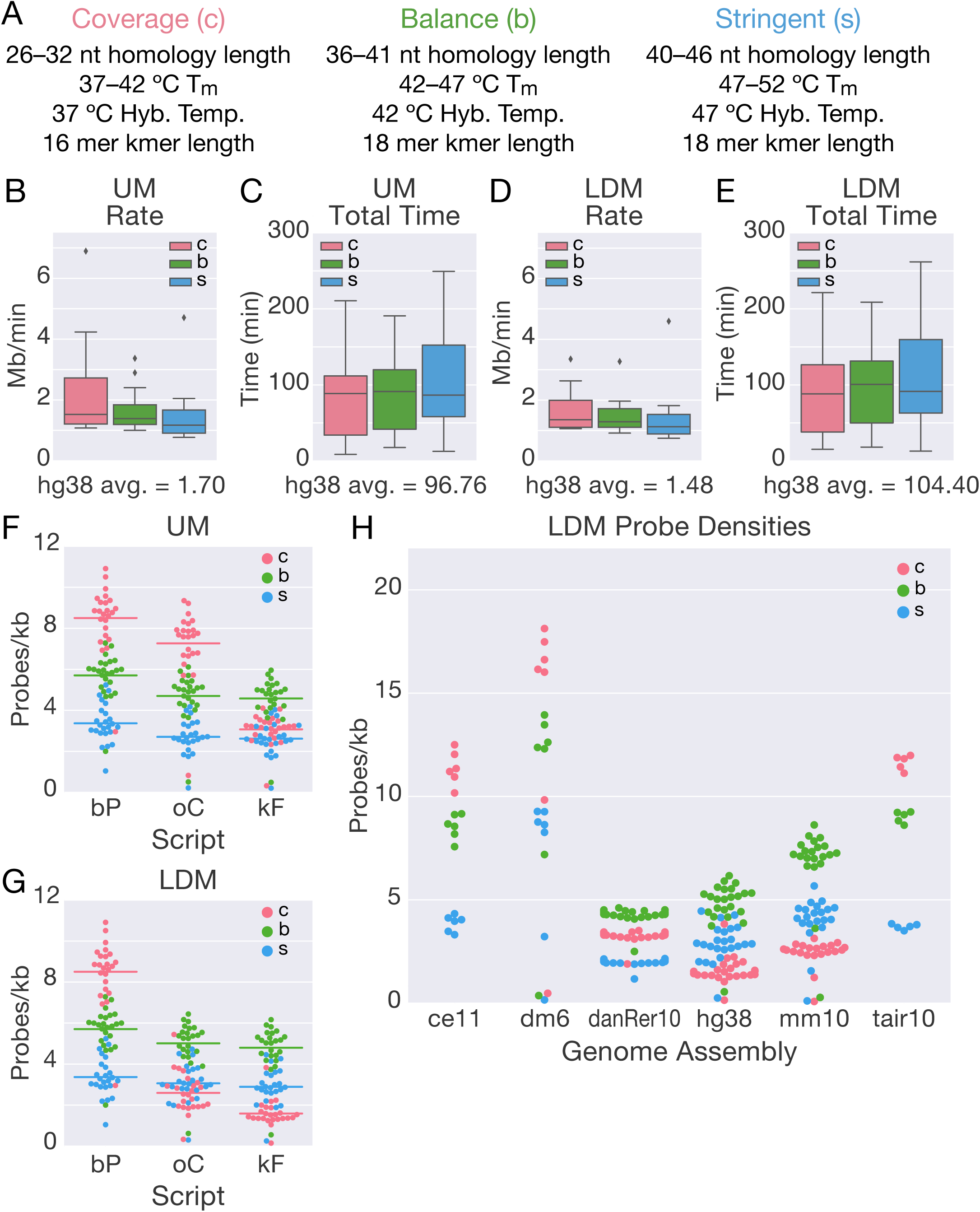
Genome-scale probe discovery with OligoMiner. (*A*) Description of three parameter sets used for genome-scale mining runs. (*B-E*) Boxplots displaying overall mining times and rates for Unique Mode (UM) (*B* and *C*) and LDA Mode (LDM) (*D* and *E*). Each chromosome was run separately and reported, resulting in 24 data points per parameter setting and a total of 72 data points per plot. The mean rate or time for all 72 data points is displayed beneath each boxplot. (*F* and *G*) Swarmplots displaying changes in probe density (i.e. probes/kb) that occurred over the course of the pipeline in (*F*) UM and (*G*) LDM. ‘bP’ = blockParse; ‘oC’ = outputClean; ‘kF’ = kmerFilter. (*H*) Swarmplot displaying probe densities in the *C. elegans* (ce11), *D. melaongaster* (dm6), zebrafish (danRer10), human (hg38), mouse (mm10), and *A. thaliana* (tair10) genome assemblies after whole-genome mining using LDM and kmerFilter.

The modularity of OligoMiner allows users to monitor how the parameters chosen at each step in the probe discovery process affect the final number of output probes. We have used this capability to examine changes in probe density (e.g. probes per kilobase) that occurred during the genome-wide probe discovery runs in hg38. As expected, blockParse discovered the highest density of candidate probes using the ‘coverage’ (c) settings, followed by ‘balance’ (b) and ‘stringent’ (s) (c: 8.5, b: 5.7, s: 3.4 probes/kb) (Fig. 2 *F* and *G*). Yet, we observed striking differences following outputClean depending on the mode used, with UM preserving the same order (c: 7.3, b: 4.7, s: 2.7 probes/kb) but the density of the ‘coverage’ oligos plummeting in LDM (c: 2.6, b: 5.0, s: 3.0 probes/kb) (Fig. 2 *F* and *G*, Note S1). We also observed large relative decreases in the density of ‘coverage’ oligos following the application of kmerFilter, but only a modest reduction with the other sets (UM c: 3.1, b: 4.6, s: 2.6 probes/kb; LDM c: 1.6, b: 4.8, s: 2.9 probes/kb) (Fig. 2 *F* and *G*); this effect is likely due to the use of 16mer dictionary with ‘coverage’ sets but an 18mer dictionary with the ‘balance’ and ‘stringent’ sets (Fig. 2*A*), a choice informed by differences in k-mer binding affinities at the different simulated hybridization temperatures (Fig. S4 and Note S1).

Collectively, our hg38 probe sets are similar in probe density to previous sets designed with Oligoarray (20, 21) and suggest that when taking the thermodynamics of hybridization into account, longer oligo probes that can support higher hybridization temperatures can effectively provide higher probe densities, as observed with the UM and LDM ‘balance’ sets (Fig. 2 *F* and *G*). Intriguingly, this phenomenon appears to depend on genome size and complexity; the same ordering of the three parameter sets was also observed in whole-genome probe discovery was performed using LDM and kmerFilter in the mouse mm10 and zebrafish danRer10 assemblies, but the ‘coverage’ set provided the highest densities in the smaller *D. melaongaster* dm6, *C. elegans* ce11, and *A. thaliana* tair10 assemblies (Fig. 2*H* and Note S1). The resulting probes discovered by these genome-scale probe discovery runs and additional LDM + kmerFilter whole-genome runs in the ce6, dm3, hg19, and mm9 assemblies will be made available on the Oligopaints website (http://genetics.med.harvard.edu/oligopaints).

### OligoMiner Enables Conventional and Super-Resolution Imaging

In order to test the efficacy of oligo probes designed with OligoMiner *in situ*, we first performed 3D FISH (54, 55) in XX 2N WI-38 human fetal lung fibroblasts with a set of 4,776 40–45mer Oligopaint probes designed using UM without kmerFilter targeting 817 kb at Xq28 (Table S1). In line with previous Oligopaint experiments using probes designed by OligoArray (20, 21), we observed highly efficient staining, with 100% of nuclei displaying at least one FISH signal and 90.3% of nuclei displaying two signals (*n* = 185; Fig. 3 *A* and *B*). We observed similarly efficient staining after performing 3D FISH in XY 2N PGP-1 fibroblasts with a set of 3,678 35–41mer Oligopaint probes designed using LDM with kmerFilter targeting 1,035 kb at 19p13.2 (Table S1) (100% nuclei with ≥1 signal, 79.2% with 2 signals, *n* = 144; Fig. 3 *B* and *C*), demonstrating the effectiveness of both the UM and LDM probe design approaches. We also validated our ability to design custom hybridization patterns by performing 3D FISH with two additional sets of 40– 45mer Oligopaint probes designed using UM without kmerFilter targeting Xq28 in WI-38 cells, which led to the expected three-color co-localization pattern *in situ* (Fig. 3*E*).

**Fig. 3.**
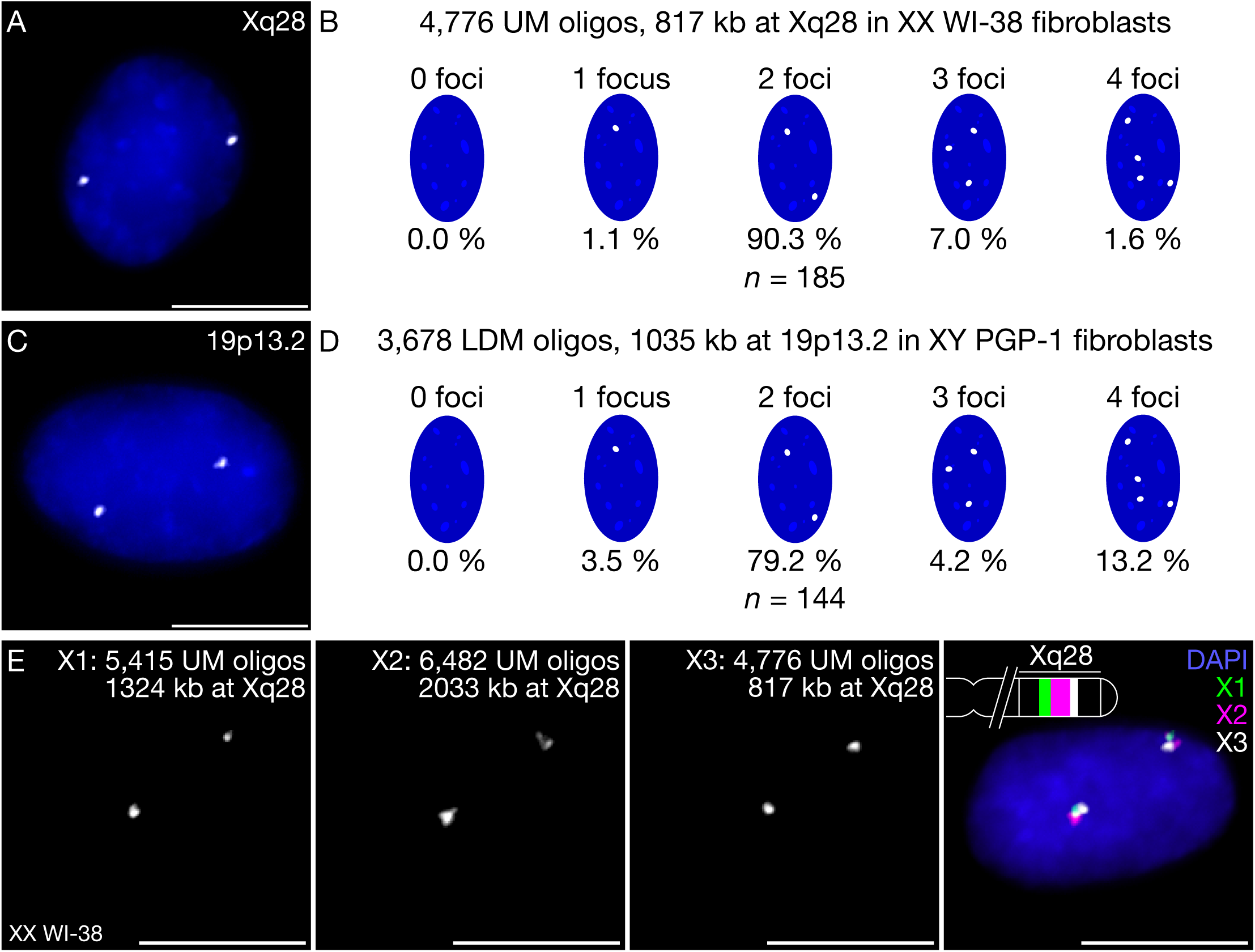
OligoMiner enables highly efficient FISH. (*A* and *B*) Representative image (*A*) and quantification (*B*) of 3D-FISH experiment performed with a probe set consisting of 4,776 UM oligos targeting 817 kb at Xq28 in human XX 2N WI-38 fibroblasts. (*C* and *D*) Representative image (*C*) and quantification (*D*) of 3D-FISH experiment performed with a probe set consisting of 3,678 LDM oligos targeting 1,035 kb at 19p13.2 in human XY 2N PGP-1 fibroblasts. (*E*) Three-color 3D FISH experiment performed using ATTO 488 labeled “X1” (green), ATTO 565 labeled “X2” (magenta), and Alexa Fluor 647 labeled “X3” UM probe sets targeting adjacent regions on Xq28 in WI-38 fibroblasts. All images in (*A*–*E*) are maximum intensity projections in Z. DNA is stained with DAPI (blue). Scale bars: 10 μm.

In order to further showcase the performance of oligos designed using OligoMiner *in situ*, we visualized 3D FISH using Stochastic Optical Reconstruction Microscopy (STORM) (56) and DNA-based Points Accumulation in Nanoscale Topography (DNA-PAINT) (57) – these single-molecule super-resolution imaging techniques spatiotemporally isolate the fluorescent emissions of individual molecules and are capable of achieving <20 nm lateral and <50 nm axial resolution, which represent an order of magnitude or more below the diffraction limit (58). Specifically, we performed STORM imaging of Oligopaints (OligoSTORM) (21) of human 19p13.2 with two sets of 35–41mer oligos designed using LDM with kmerFilter targeting either a 1,035 kb region with 3,768 oligos (Table S1) (Fig. 4 *A* and *B*) or a 20 kb region with 104 oligos (Table S1) (Fig. 4 *C* and *D*) and in both cases were readily able to resolve the nanoscale morphologies of these foci, including features <40 nm (Fig. 4*D*), values comparable to those obtained using probes designed by OligoArray (21). We also performed DNA-PAINT imaging of Oligopaints (OligoDNA-PAINT) (21) to visualize our 817 kb Xq28 probe set (Table S1) (Fig. 4 *E* and *F*) and a set of 167 35–41mer oligos designed using LDM with kmerFilter targeting the Xist RNA (59) (Table S1) (Fig. 4 *G* and *H*), which also enabled us to reveal <40 nm structural features in the super-resolved images (Fig. 4 *F* and *H*). Taken together, these super-resolution experiments demonstrate the OligoMiner oligos can readily enable the single-molecule super-resolution imaging of a broad range of target types and sizes.

**Fig. 4.**
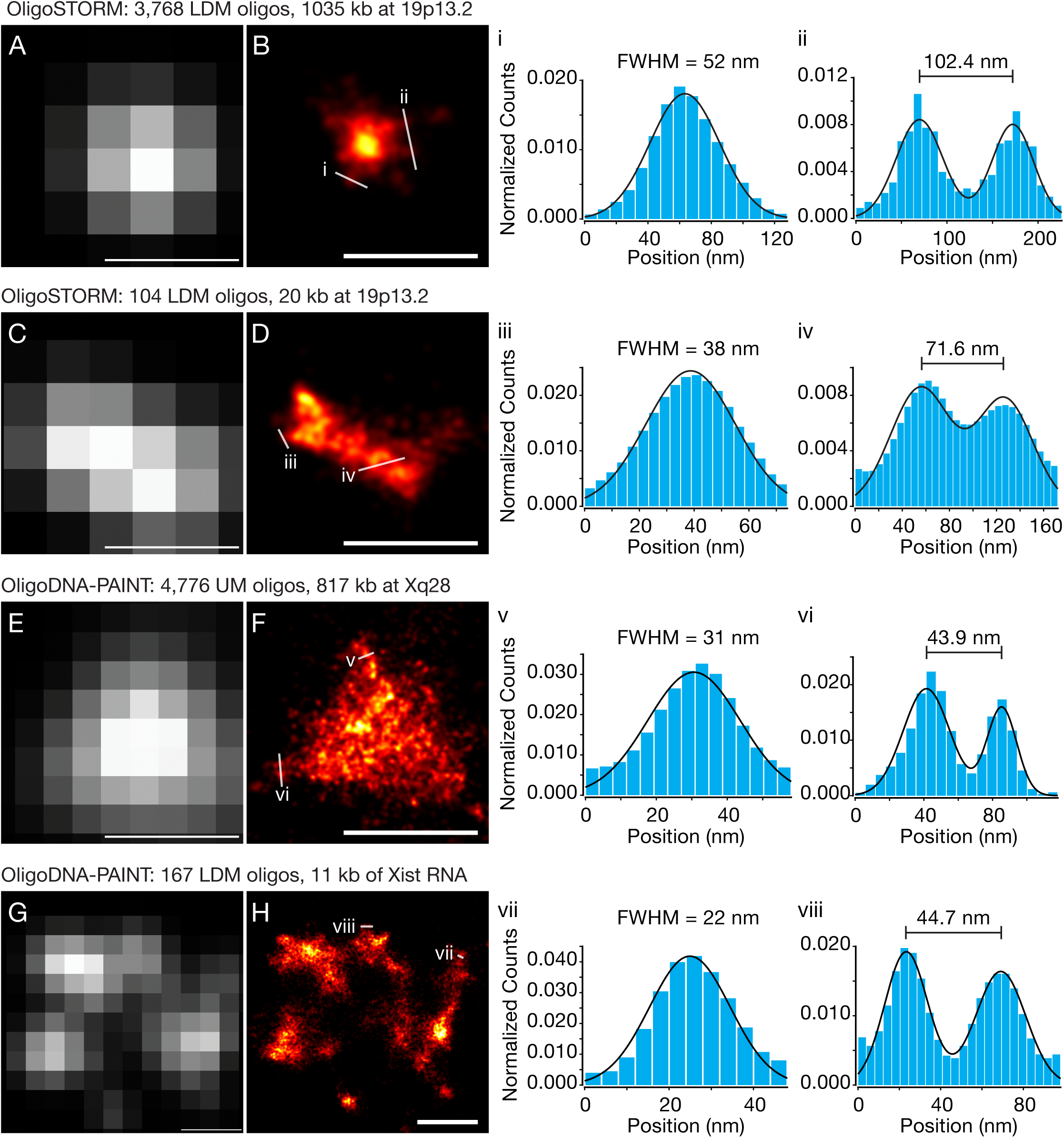
Single-molecule super-resolution imaging of OligoMiner oligos. (*A* and *B*) Diffraction-limited (*A*) and super-resolved STORM (*B*) images of a probe set consisting of 3,678 LDM oligos targeting 1,035 kb at 19p13.2 in human XY 2N PGP-1 fibroblasts. (*C* and *D*) Diffraction-limited (*C*) and super-resolved STORM (*D*) images of a probe set consisting of 104 LDM oligos targeting 20 kb at 19p13.2 in PGP-1 fibroblasts. (*E* and *F*) Diffraction-limited (*E*) and super-resolved DNA-PAINT (*F*) images of a probe set consisting of 4,776 UM oligos targeting 817 kb at Xq28 in human XY 2N MRC-5 fibroblasts. (*G* and *H*) Diffraction-limited (*G*) and super-resolved DNA-PAINT (*H*) images of a probe set consisting of 176 LDM oligos targeting 11 kb of the Xist RNA in human XX 2N WI-38 fibroblasts. (*i*–*viii*) present normalized single-molecule counts along the indicated 1-dimensional line traces (blue bars) and 1- or 2-component Gaussian fits to the underyling data (black lines). Super-resolution data is presented using a ‘Hot’ colormap in which single-molecule localization density scales from black (lowest) to red to yellow to white (highest). Scale bars: 500 nm.

## Discussion

OligoMiner provides a framework for the rapid design of oligo hybridization probes on the genome-wide scale. We have demonstrated the ease and scalability of our pipeline by mining the human hg38 genome assembly with three distinct parameter sets and in two specificity-checking modes, a feat that would have otherwise required many months of cluster computing, and further highlighted the effectiveness of our approach with conventional and single-molecule super-resolution imaging. Written in open source Python and Biopython and freely available via GitHub (https://github.com/brianbeliveau/OligoMiner), OligoMiner can readily be run on any standard laptop or desktop computer and exclusively uses standard bioinformatic file formats, providing users the opportunity to integrate OligoMiner scripts into existing pipelines and readily allowing additional and updated programs to be seamlessly integrated into the workflow. Critically, OligoMiner is capable of discovering the thousands to tens of thousands of oligo probes commonly ordered as pools from commercial suppliers in mere minutes, freeing the researcher to tailor the design of each probe set to the experimental question at hand instead of relying on pre-existing collections of probe sequences obtained from previous probe mining runs or online databases (20). Accordingly, we anticipate that OligoMiner could be employed more broadly to design hybridization probes for a wide range of experimental assays beyond *in situ* hybridization.

## Methods

### Genome Sequences

The hg19, hg38, mm9, mm10, ce6, ce11, danRer10, dm3, and dm6 genome assemblies were downloaded both with and without repeat masking from http://genome.ucsc.edu. The tair10 assembly was downloaded from http://arabidopsis.org. To generate a repeat-masked version of tair10, transposable element locations identified by TASR (60) were converted to BED format and used as a guide for masking by pyfaidx (61).

### Pipeline Construction and Implementation

OligoMiner is written for Python 2.7 and depends on Biopython (35) and scikit-learn 0.17+ (50). Additional optional dependencies include Jellyfish 2.0+ (51) for k-mer screening and NUPACK 3.0 (47–49) for secondary structure analysis. To generate data for this study, scripts were either executed locally in an OS X Anaconda Python 2.7 environment (Continuum Analytics) created with the command ‘conda create --name probeMining biopython scikit-learn’ or in a CentOS Linux environment on the Orchestra High Performance Compute Cluster at Harvard Medical School.

### LDA Model Construction

Two sets of ‘probe’ and ‘target site’ sequences were used for the LDA model construction. For the first, all possible k-mers ≥8 were generated from 500 40–46mer sequences from hg38 chrX that were identified as candidate probes by blockParse, resulting in a total pool of 337,514 truncated and full-length sequences. In the second, 100 26–32mer, 100 35–41mer, and 100 40– 46mer sequences from hg38 chr7 identified as candidate probes by blockParse were used as a starting pool of sequences. A Python script was then used to generate variant sequences containing 1–10 point mutations, 1–3 insertions of 1–6 bases each, or 1–3 deletions of 1–6 bases each, resulting in a total pool of 69,300 parental and variant sequences. These two pools were then combined to create a final pool of 406,814 sequences. In order to generate Bowtie2 alignment scores for each ‘probe’ – ‘target-site’ pairing, the ‘probe’ sequence flanked by 3 ‘T’ bases on both the 5’ and 3’ ends was used to create a Bowtie2 alignment index, against which the ‘target-site’ sequence was aligned using the settings: ‘--local -D 20 -R 3 -N 1 -L 10 -i S,1,0.5 --score-min G,1,1 -k 1.’ To generate NUPACK duplexing probabilities for each pairing at a given temperature, the ‘complexes’ executable was first called and given an input of the reverse complement of the ‘probe’ sequence flanked by 3 ‘T’ bases on both the 5’ and 3’ ends and the ‘target-site’ sequence in a two-strand simulation with a maximum complex size of two strands. To account for FISH conditions, the Na^+^ concentration was set to 390 mM and the input temperature was increased by 31°C (0.62 * 50) to account for the presence of 50% formamide. The resulting partition function outputted by ‘complexes’ was then passed to the ‘concentrations’ executable, with each strand being assigned an initial concentration of 1 μM. The percentage of the ‘probe’ oligo contained in the ‘probe-target’ complex was then stored as the duplexing probability. If the probability of duplexing was <0.2, the pairing was assigned to the ‘not likely to bind stably’ / (-1) class; If the probability of duplexing was ≥ 0.2, the pairing wa assigned to the ‘likely to bind stably’ / (1) class. LDA model building, testing, and validation was performed using scikit-learn 0.17 (50).

### Whole-Genome Probe Discovery

Genome assemblies in FASTA format without repeat masking were used to build Bowtie2 alignment indices and Jellyfish files. Repeat-masked input files were used for probe discovery. The blockParse script was run with the settings indicated in Figure 2*A* and all other values set to their defaults (Fig. S1). Bowtie2 was run with ‘--very-sensitive-local –k 2 -t’ in UM and ‘--local -D 20 -R 3 -N 1 -L 20 -i C,4 --score-min G,1,4 -k 2 -t’ in LDM. The outputClean script was run with default values (Fig. S2) in either LDM or UM. The kmerFilter script was used with the k-mer lengths indicated in Figure 2*A* and ‘-k/--kmerThreshold’ set to 5. To minimize file sizes and maximize speed, Jellyfish files were created such that k-mers occurring 0 or 1 times were not recorded and all kmers occurring >255 times were reported as ‘255’, i.e. the counts were recorded with 1 bit. Jellyfish hash size was set to roughly the size of the genome assembly. E.g. the command ‘jellyfish count -s 3300M -m 18 -o hg38_18.jf --out-counter-len 1 -L 2 hg38.fa’ was used to create the 18mer dictionary for hg38. Bowtie 2.2.4 and Jellyfish 2.2.4 were used. The resulting probe files for all whole-genome runs described in Figure 2 as well as whole-genome runs with the ‘c’, ‘b’, and ‘s’ parameter sets in the hg19, mm9, dm3, and ce6 assemblies using LDM and kmerFilter will be made available at http://genetics.med.harvard.edu/oligopaints.

### Mining Speed Calculations

Genome-scale hg38 mining runs were conducted on the Orchestra Compute cluster, with each chromosome being run as its own individual job (i.e. without further parallelization) for each step in the probe design process (blockParse, Bowtie2, outputClean, kmerFilter). Wall clock times for the three OligoMiner Python scripts were reported via the Python ‘timeit’ module and written to meta files by flagging the ‘-M/--Meta’ option present in the three scripts. Bowtie2 wall clock time was reported by flagging the ‘-t’ option and read from the printed output. Graphs presenting probe mining speed and probe densities were created in Python using seaborn (62).

### Oligopaint Probe Synthesis

For the OligoMiner settings used to design each Oligopaint FISH probe set, please see Table S1. Probe sets were synthesized using the previously described gel extraction (20) (Xq28 probes) or T7 methods (24) (19p13.2 probes) and generated from complex oligo libraries ordered from Custom Array (Bothell, WA). For a stepwise synthesis protocol, please see (63). The Xist RNA FISH probe set was ordered as a set of individually column synthesized oligos from Integrated DNA Technologies (Coralville, IA). For a list of primer sequences used, please see Table S2.

### Cell Culture

Human WI-38 (ATCC CCL-75), MRC-5 (ATCC CCL-171), and PGP-1 fibroblasts (Coriell Institute GM23248) were grown at 37°C in the presence of 5% CO_2_ in Dulbecco’s Modified Eagle Medium (Gibco 10564) supplemented with 10% (vol/vol) serum (Gibco 10437), 50□U/ml penicillin, and 50□μg/ml streptomycin (Gibco 15070). The PGP-1 fibroblasts were also supplemented with MEM non-essential amino acids solution (Gibco 11140050).

### 3D DNA FISH

3D DNA FISH (54, 55) was essentially performed as described previously (20–22, 63). WI-38, IMR-90, or PGP-1 fibroblasts were seeded at ∼20% confluence into the wells of Labtek-II Coverglass Chambers, ididi coverglass chambers, or onto #1.5 coverglass and allowed to grow to ∼70-90% confluence in a mammalian tissue culture incubator. Samples were then rinsed with 1x PBS and fixed for 10 min in 1x PBS + 4% (wt/vol) paraformaldehyde, then rinsed again with 1x PBS. Samples were next permeabilized by a rinse in 1x PBS + 0.1% (vol/vol) Tween-20, then a 10 min incubation in 1x PBS + 0.5% (vol/vol) Triton X-100, then a 5 minute incubation in 0.1 N HCl. Samples were then transferred to 2x SSC + 0.1% (vol/vol) Tween-20 (SSCT), then to 2x SSCT + 50% (vol/vol) formamide. Samples were then incubated in 2x SSCT + 50% formamide at 60°C for 20–60 min, after which a hybridization solution consisting of 2x SSCT, 50% formamide, 10% (wt/vol) dextran sulfate, 40 ng/μl RNase A (Thermo Fisher EN0531), and Oligopaint FISH probe sets at 1.6 or 2.5 μM was added. Samples were denatured at 78°C for 3 minutes on a water-immersed heat block or flat-block thermocycler (Eppendorf Mastercycler Nexus) and then allowed to hybridize for 24+ hours at 47–52°C either in a humidified chamber placed in an air incubator or on a flat-block thermocycler. After hybridization, samples were washed in 2x SSCT at 60°C for 5 minutes four times, then 2x SSCT at room temperature two times, then transferred to 1x PBS. Unlabeled secondary oligos (21) and tertiary oligos bearing Alexa Fluor 405 and 647 dyes (22) (Table S2) at 0.5–1 μM were subsequently hybridized to the 19p13.2 samples for 1 hour in 2x SSC + 30% formamide + 10% dextran sulfate at room temperature and washed three times for 5 min each in 2x SSC + 30% formamide. SlowFade Gold + 4’,6-diamidino-2-phenylindole (DAPI) (ThermoFisher S36938) was added to samples prepared for diffraction-limited imaging. Samples for super-resolution imaging were stained in a 1 μg/ml DAPI solution in 1x PBS or 2x SSCT for 5 min at 37°C, followed by a brief rinse in 1x PBS or 2x SSCT at room temperature.

### RNA FISH

RNA FISH was performed exactly as described for ‘3D DNA FISH’, except that the 3 min denaturation at 78°C was replaced with a 5 min incubation at 60°C, RNase A was omitted from the hybridization buffer, and hybridization was carried out at 42°C for 16 hours.

### Diffraction-Limited Imaging and Quantification of FISH Efficiency

Diffraction-limited imaging of 3D DNA FISH samples was conducted on an inverted Zeiss Axio Observer Z1 using a 63x Plan-Apochromat Oil DIC (N.A. 1.40) objective. Samples were illuminated using Colibri light source using a 365 nm, 470 nm, 555 nm, or 625 nm LED. DAPI was visualized using a filter set composed of a 365 nm clean-up filter (G 365), a 395 nm long-pass dichroic mirror (FT 395), and a 445/50 nm bandpass emission filter (BP 445/50). ATTO 488 was visualized using a filter set composed of a 470/40 nm excitation filter (BP 470/40), a 495 nm long-pass dichroic mirror (FT 495), and a 525/50 nm bandpass emission filter (BP 525/50). ATTO 565 was visualized using a filter set composed of a 545/25 nm excitation filter (BP 545/25), a 570 nm long-pass dichroic mirror (FT 570), and a 605/70 nm bandpass emission filter (BP 605/70). Alexa Fluor 647 was visualized using a filter set composed of a 640/30 nm excitation filter (BP 640/30), a 660 nm long-pass dichroic mirror (FT 660), and a 690/50 nm bandpass emission filter (BP 690/50). Images were acquired using a Hamamatsu Orca-Flash 4.0 sCMOS camera with 6.5 μm pixels, resulting in an effective magnified pixel size of 103 nm. Z-stacks were acquired using an interval of 240 nm. Images were processed using Zeiss Zen software and Fiji/ImageJ (64). FISH foci were identified manually by scanning through Z-stacks; signals whose center-to-center distance was <1 μm were considered to be a single focus.

### STORM Imaging

STORM imaging was performed on a commercial Nikon N-STORM 3.0 microscope featuring a Perfect Focus System and a motorized TIRF illuminator at the Nikon Imaging Center located at Harvard Medical School. STORM was performed using highly inclined and laminated optical sheet illumination (HILO) (65) and with pulsed activation of the 405 nm laser, followed by 647 nm, and then 561. Light was focused through a CFI Apo TIRF 100x Oil (N.A. 1.49) objective. The 561 nm laser was used at 2% (out of 50 mW) to image 200 nm orange FluoSpheres (F8809, ThermoFisher), which were used as fiducial makers to facilitate drift correction. The 405 nm laser was used to enhance the blinking rate at 0–5% (out of 20 mW), and the 647 nm laser was used at 100% power (out of 125 mW measured at fiber optic). Emission light was spectrally filtered (Chroma ET600/50m for 561; Chroma ET700/75m for 647) and imaged on an EMCCD camera (Andor iXon X3 DU-897) with 16 μm pixels using a CCD readout bandwidth of 10 MHz at 16□bit, 1 bit pre-amp gain and no electron-multiplying gain on the center 256 x 256 or 186 x 190 pixels, resulting in an effective pixel size of 160 nm. 6,250 or 12,500 10 ms frames were acquired. Single-molecule localization events were identified using in-house MATLAB software (66) that calls a 2D fitting algorithm (67). Individual localization events were blurred with 2D Gaussian functions whose ‘sigma’ parameter was set according to the global drift-independent localization precision as determined by Nearest Neighbor based Analysis (NeNA) (68). NeNA values: 19p13.2 1035 kb– 12.6 nm sigma, 29.6 nm supported resolution; 19p13.2 20 kb– 11.8 nm sigma, 27.6 nm supported resolution. 1-and 2-component Gaussian fits of the line traces presented in Figure 4*A*–*D* were calculated using the ‘Gaussian Mixture Model’ module in scikit-learn (50).

### DNA-PAINT Imaging

DNA-PAINT imaging was performed on a commercial Nikon N-STORM 3.0 microscope featuring a Perfect Focus System and a motorized TIRF illuminator. DNA-PAINT was performed using HILO with a 15-30% of a 200 mW 561 nm laser (Coherent Sapphire) using a CFI Apo TIRF 100x Oil (N.A. 1.49) objective at an effective power density of ∼0.5–1 kW/cm^2^. 561 nm laser excitation light was passed through a clean-up filter (Chroma ZET561/10) and directed to the objective using a multi-band beam splitter (Chroma ZT405/488/561/647rpc). Emission light was spectrally filtered (Chroma ET600/50m) and imaged on an EMCCD camera (Andor iXon X3 DU-897) with 16 μm pixels using a CCD readout bandwidth of 3 MHz at 14□bit, 5.1 pre-amp gain and no electron-multiplying gain on the center 256 x 256 pixels, resulting in an effective pixel size of 160 nm. 15,000 100 ms frames were acquired for each image using 1–3 nM of Cy3B-labeled 10mer oligo in 1x PBS + 125–500 nM NaCl. 40 nm gold nanoparticles (Sigma-Aldrich 753637) were used as fiducial markers to facilitate drift correction. Single-molecule localization events were identified using in-house MATLAB software (66) that calls a 2D fitting algorithm (67). Individual localization events were blurred with 2D Gaussian functions whose ‘sigma’ parameter was set according to the global drift-independent localization precision as determined by NeNA (68). NeNA values: Xq28– 5.6 nm sigma, 13.2 nm supported resolution; Xist RNA– 5.1 nm sigma, 12.0 nm supported resolution. 1- and 2-component Gaussian fits of the line traces presented in Figure 4*E*–*H* were calculated using the ‘Gaussian Mixture Model’ module in scikit-learn (50).

## Acknowledgements

We thank Ninning Liu, Mingjie Dai, Thomas C. Ferrante, Josh Rosenberg, Nikhil Gopalkrishnan, Florian Schüder, Ralf, Jungmann, and members of the Yin and Wu labs for helpful discussions, Jin Billy Li for the idea to use k-mer filtering as a means of specificity checking, and Geoffrey Fudenberg for assistance with the 19p13.2 1,035 kb and 20 kb probe design. This work was supported by awards to PY from the NIH (NIH 1R01EB018659-01 and NIH 1-U01-MH106011-01), ONR (N00014-13-1-0593, N00014-14-1-0610, N00014-16-1-2182, N00014-16-1-2410), and NSF (CCF-1317291) and awards to CTW from the NIH (DP1GM106412, R01HD091797). BJB was supported by a Damon Runyon Cancer Research Foundation Fellowship. HMS was supported by a Uehara Memorial Foundation Research Fellowship. SKS was supported by Postdoctoral Fellowships from EMBO and the Human Frontier Science Program. JYK and SCN were supported by NSF Graduate Research Fellowships.

---

**Fig. S1.** Description of blockParse. (*A*) Schematic diagram illustrating the nature and order of checks used to screen candidate probe sequences. (*B*) Description of command-line options and the default values/settings for each.

**Fig. S2.** Description of outputClean. (*A*) Schematic diagram illustrating the task order of the script. (*B*) Description of command-line options and the default values/settings for each.

**Fig. S3.** Summary information for each temperature-specific LDA model. (*A*–*F*) For each temperature, the Precision, Recall, support-weighted F_1_ Score, and Support are given.

**Fig. S4.** Boxplots depicting the duplexing probabilities of all kmers ≥8 nucleotides (nt) in length with the reverse complements of their 40–46mer parental seqeunces at six different simulation temperatures in 390 mM Na^+^ and 50% formamide.

**Table S1.** Description of Oligopaint probe sets used.

**Table S2.** Description of oligo sequences used for probe set generation and visualization.

## References

1. Pardue ML, Gall JG (1969) Molecular hybridization of radioactive DNA to the DNA of cytological preparations. Proc Natl Acad Sci U S A 64(2):600–4.

2. John HA, Birnstiel ML, Jones KW (1969) RNA-DNA hybrids at the cytological level. Nat Publ Gr 223:582–587.

3. Buongiorno-Nardelli M, Amaldi F (1970) Autoradiographic Detection of Molecular Hybrids between rRNA and DNA in Tissue Sections. Nature 225(5236):946–948.

4. Lawrence JB, Singer RH (1985) Quantitative analysis of in situ hybridization methods for the detection of actin gene expression. Nucleic Acids Res 13(5):1777–99.

5. van der Ploeg M (2000) Cytochemical nucleic acid research during the twentieth century. Eur J Histochem 44(1):7–42.

6. Levsky JM, Singer RH (2003) Fluorescence in situ hybridization: past, present and future. J Cell Sci 116(14):2833–2838.

7. Riegel M (2014) Human molecular cytogenetics: From cells to nucleotides. Genet Mol Biol 37(1 SUPPL. 1):194–209.

8. Rigby PWJ, Dieckmann M, Rhodes C, Berg P (1977) Labeling deoxyribonucleic acid to high specific activity in vitro by nick translation with DNA polymerase I. J Mol Biol 113(1):237–251.

9. Langer PR, Waldrop AA, Ward DC (1981) Enzymatic synthesis of biotin-labeled polynucleotides: novel nucleic acid affinity probes. Proc Natl Acad Sci U S A 78(11):6633–6637.

10. Landegent JE, Jansen in de Wal N, Dirks RW, van der Ploeg M (1987) Use of whole cosmid cloned genomic sequences for chromosomal localization by non-radioactive in situ hybridization. Hum Genet 77(4):366–370.

11. Moyzis RK, et al. (1988) A highly conserved repetitive DNA sequence, (TTAGGG)n, present at the telomeres of human chromosomes. Proc Natl Acad Sci U S A 85(18):6622–6.

12. Matera A, Ward D (1992) Oligonucleotide probes for the analysis of specific repetitive DNA sequences by fluorescence in situ hybridization. Hum Mol Genet 1(7):535–539.

13. Dernburg AF, et al. (1996) Perturbation of nuclear architecture by long-distance chromosome interactions. Cell 85(5):745–759.

14. Dirks RW, et al. (1990) Simultaneous detection of different mRNA sequences coding for neuropeptide hormones by double in situ hybridization using FITC- and biotin-labeled oligonucleotides. J Histochem Cytochem 38(4):467–473.

15. Femino a M, Fay FS, Fogarty K, Singer RH (1998) Visualization of single RNA transcripts in situ. Science 280(5363):585–590.

16. Raj A, van den Bogaard P, Rifkin S a, van Oudenaarden A, Tyagi S (2008) Imaging individual mRNA molecules using multiple singly labeled probes. Nat Methods 5(10):877–879.

17. Kosuri S, Church GM (2014) Large-scale de novo DNA synthesis: technologies and applications. Nat Methods 11(5):499–507.

18. Yamada NA, et al. (2011) Visualization of fine-scale genomic structure by oligonucleotide-based high-resolution FISH. Cytogenet Genome Res 132(4):248–254.

19. Boyle S, Rodesch MJ, Halvensleben HA, Jeddeloh JA, Bickmore WA (2011) Fluorescence in situ hybridization with high-complexity repeat-free oligonucleotide probes generated by massively parallel synthesis. Chromosom Res 19(7):901–909.

20. Beliveau BJ, et al. (2012) Versatile design and synthesis platform for visualizing genomes with Oligopaint FISH probes. Proc Natl Acad Sci 109(52):21301–21306.

21. Beliveau BJ, et al. (2015) Single-molecule super-resolution imaging of chromosomes and in situ haplotype visualization using Oligopaint FISH probes. Nat Commun 6(May):7147.

22. Boettiger AN, et al. (2016) Super-resolution imaging reveals distinct chromatin folding for different epigenetic states. Nature 529(7586):418–22.

23. Wang S, et al. (2016) Spatial organization of chromatin domains and compartments in single chromosomes. Science (80- ) 353(6299):598–602.

24. Chen KH, Boettiger AN, Moffitt JR, Wang S, Zhuang X (2015) RNA imaging. Spatially resolved, highly multiplexed RNA profiling in single cells. Science 348(6233):aaa6090.

25. Shah S, Lubeck E, Zhou W, Cai L (2016) In Situ Transcription Profiling of Single Cells Reveals Spatial Organization of Cells in the Mouse Hippocampus. Neuron 92(2):342–357.

26. Pernthaler J, Glöckner FO, Schönhuber W, Amann R (2001) Fluorescence in situ hybridization with rRNA-targeted oligonucleotide probes. Methods Microbiol 30(3):207–226.

27. Yilmaz LS, Parnerkar S, Noguera DR (2011) MathFISH, a web tool that uses thermodynamics-based mathematical models for in silico evaluation of oligonucleotide probes for fluorescence in situ hybridization. Appl Environ Microbiol 77(3):1118–1122.

28. Rogan PK, Cazcarro PM, Knoll JHM (2001) Sequence-Based Design of Single-Copy Genomic DNA Probes for Fluorescence In Situ Hybridization Sequence-Based Design of Single-Copy Genomic DNA Probes for Fluorescence In Situ Hybridization. 1086–1094.

29. Navin N, et al. (2006) PROBER: Oligonucleotide FISH probe design software. Bioinformatics 22(19):2437–2438.

30. Nedbal J, Hobson PS, Fear DJ, Heintzmann R, Gould HJ (2012) Comprehensive FISH Probe Design Tool Applied to Imaging Human Immunoglobulin Class Switch Recombination. PLoS One 7(12). doi:10.1371/journal.pone.0051675.

31. Bienko M, et al. (2013) A versatile genome-scale PCR-based pipeline for high-definition DNA FISH. Nat Methods 10(2):122–4.

32. Banér J, et al. (2003) Parallel gene analysis with allele-specific padlock probes and tag microarrays. Nucleic Acids Res 31(17):e103.

33. Stenberg J, Nilsson M, Landegren U (2005) ProbeMaker: an extensible framework for design of sets of oligonucleotide probes. BMC Bioinformatics 6:229.

34. Rouillard JM, Zuker M, Gulari E (2003) OligoArray 2.0: Design of oligonucleotide probes for DNA microarrays using a thermodynamic approach. Nucleic Acids Res 31(12):3057–3062.

35. Cock PJA, et al. (2009) Biopython: Freely available Python tools for computational molecular biology and bioinformatics. Bioinformatics 25(11):1422–1423.

36. Lipman DJ, Pearson WR (1985) Rapid and sensitive protein similarity searches. Science (80-) 227(4693):1435–1441.

37. Smit A, Hubley R, Green P (2013) RepeatMasker Open-4.0. 2013&2015. http://www.repeatmasker.org. Available at: http://repeatmasker.org.

38. SantaLucia J (1998) A unified view of polymer, dumbbell, and oligonucleotide DNA nearest-neighbor thermodynamics. Proc Natl Acad Sci U S A 95(4):1460–5.

39. Cock PJA, Fields CJ, Goto N, Heuer ML, Rice PM (2010) The Sanger FASTQ file format for sequences with quality scores, and the Solexa/Illumina FASTQ variants. Nucleic Acids Res 38(6):1767–1771.

40. Langmead B, Trapnell C, Pop M, Salzberg SL (2009) Bowtie: An ultrafast memory-efficient short read aligner. [http://bowtie.cbcb.umd.edu/]. Genome Biol (10):pR25.

41. Langmead B, Salzberg SL (2012) Fast gapped-read alignment with Bowtie 2. Nat Methods 9(4):357–359.

42. Li H, Durbin R (2010) Fast and accurate long-read alignment with Burrows-Wheeler transform. Bioinformatics 26(5):589–595.

43. Altschul SF, Gish W, Miller W, Myers EW, Lipman DJ (1990)Altschul et al.. 1990. Basic Local Alignment Search Tool. J Mol Biol 215(3):403–410.

44. Li H, et al. (2009) The Sequence Alignment/Map format and SAMtools. Bioinformatics 25(16):2078–2079.

45. Kent WJ, et al. (2002) The Human Genome Browser at UCSC. Genome Res 12(6):996–1006.

46. Quinlan AR, Hall IM (2010) BEDTools: A flexible suite of utilities for comparing genomic features. Bioinformatics 26(6):841–842.

47. Dirks RM, Pierce NA (2003) A partition function algorithm for nucleic acid secondary structure including pseudoknots. J Comput Chem 24(13):1664–1677.

48. Dirks RM, Pierce NA (2004) An algorithm for computing nucleic acid base-pairing probabilities including pseudoknots. J Comput Chem 25(10):1295–1304.

49. Dirks RM, Bois JS, Schaeffer JM, Winfree E, Pierce NA (2007) Thermodynamic Analysis of Interacting Nucleic Acid Strands. SIAM Rev 49(1):65–88.

50. Pedregosa F, Varoquaux G (2011) Scikit-learn: Machine learning in Python doi:10.1007/s13398-014-0173-7.2.

51. Marçais G, Kingsford C (2011) A fast, lock-free approach for efficient parallel counting of occurrences of k-mers. Bioinformatics 27(6):764–770.

52. Arvey A, et al. (2010) Minimizing off-target signals in RNA fluorescent in situ hybridization. Nucleic Acids Res 38(10). doi:10.1093/nar/gkq042.

53. Moffitt JR, et al. (2016) High-throughput single-cell gene-expression profiling with multiplexed error-robust fluorescence in situ hybridization. Proc Natl Acad Sci U S A 113(39):11046–51.

54. Solovei I, et al. (2002) Spatial preservation of nuclear chromatin architecture during three-dimensional fluorescence in situ hybridization (3D-FISH). Exp Cell Res 276(1):10–23.

55. Solovei I, Cremer M (2010) 3D-FISH on cultured cells combined with immunostaining. Methods Mol Biol 659:117–126.

56. Rust MJ, Bates M, Zhuang XW (2006) Sub-diffraction-limit imaging by stochastic optical reconstruction microscopy (STORM). Nat Methods 3(10):793–795.

57. Jungmann R, et al. (2010) Single-molecule kinetics and super-resolution microscopy by fluorescence imaging of transient binding on DNA origami. Nano Lett 10(11):4756–4761.

58. Godin AG, Lounis B, Cognet L (2014) Super-resolution microscopy approaches for live cell imaging. Biophys J 107(8):1777–1784.

59. Brown CJ, et al. (1992) The human XIST gene: Analysis of a 17 kb inactive X-specific RNA that contains conserved repeats and is highly localized within the nucleus. Cell 71(3):527–542.

60. El Baidouri M, et al. (2015) A new approach for annotation of transposable elements using small RNA mapping. Nucleic Acids Res 43(13):e84–e84.

61. Shirley M, Ma Z, Pedersen B, Wheelan S (2015) Efficient “pythonic” access to FASTA file using pyfaidx. PeerJ Prepr:1–4.

62. Waskom M, et al. (2014) seaborn: v0.5.0 (November 2014). doi:10.5281/ZENODO.12710.

63. Beliveau BJ, Apostolopoulos N, Wu C ting (2014) Visualizing genomes with Oligopaint, fish probes. Curr Protoc Mol Biol 2014(January):14.23.1–14.23.20.

64. Schindelin J, et al. (2012) Fiji: an open-source platform for biological-image analysis. Nat Methods 9(7):676–682.

65. Tokunaga M, Imamoto N, Sakata-Sogawa K (2008) Highly inclined thin illumination enables clear single-molecule imaging in cells. Nat Methods 5(2):159–61.

66. Dai M, Jungmann R, Yin P (2016) Optical imaging of individual biomolecules in densely packed clusters. Nat Nanotechnol 11(9):798–807.

67. Smith CS, Joseph N, Rieger B, Lidke KA (2010) Fast, single-molecule localization that achieves theoretically minimum uncertainty. Nat Methods 7(5):373–375.

68. Endesfelder U, Malkusch S, Fricke F, Heilemann M (2014) A simple method to estimate the average localization precision of a single-molecule localization microscopy experiment. Histochem Cell Biol 141(6):629–638.

